# Building an ‘epigenetic clock’: Utilizing whole genome DNA methylation patterns to predict age in haddock, *Melanogrammus aeglefinus*

**DOI:** 10.1101/2025.10.16.682892

**Authors:** E.L. Strand, R. McBride, E. Robillard, A.G. Bodnar, S.A. Wanamaker, T.P. O’Donnell

**Author notes:** Corresponding author Emma L. Strand.

## Abstract

Age-based population models are a gold standard approach to estimate stock size and sustainable catch recommendations for effective fisheries management. Traditional ageing by counting annuli on their ear stones (otoliths), is labor-intensive, lethal, and can be inaccurate for long-lived fishes. In this study, we present an accurate, non-lethal epigenetic clock method for predicting age in haddock, *Melanogrammus aeglefinus*, one of the most valuable commercial fisheries in the north Atlantic Ocean. Pectoral fin tissue (n=140 individuals) was collected during Fall 2022 and Spring 2023 bottom trawl surveys in the Georges Bank-Gulf of Maine region, and whole-genome bisulfite sequencing was conducted to measure CpG loci DNA methylation levels. The most age-predictive loci were identified using a three-step modeling approach: a Bayesian generalized linear mixed model with length and sex as random factors followed by machine learning models with a 70/30 train-test split. The final model used 111 loci to predict age within 1-year at 93% accuracy. Hypo- and hyper-methylated age-related loci were found throughout the genome and were enriched in intragenic regions with functions related to immunity, transcriptional regulation, and signal transduction. This study provides an accurate, cost-effective, and scalable epigenetic tool for fisheries management to predict the age of haddock.

## Introduction

Demographic assessments of populations are crucial for the sustainable management of fish stocks. Models that require age structure information provide the most accurate estimates of population dynamics because they can estimate stock development temporally, while also calculating uncertainty estimates^1^. Unreliable age information can lead to unsustainable fishing policies or conversely, unrealized economic value for fisheries worldwide^2,3^, as shown by several historical examples. Populations of orange roughy (*Hoplostethus atlanticus*; Trachichthyidae) off New Zealand faced near collapse due to an underestimated lifespan of 20-30 years, later corrected to 150 years^4,5^. Additionally, several species of Sebastes rockfishes experienced population decline from underestimating ages of species that can live over 200 years^6,7^ and walleye pollock *(Gadus chalcogrammus*, Gadidae) in the Bering Sea saw a 99% decline in commercial catch in less than a decade due to ageing errors^8,9^. Well-managed fisheries rely on age information, which doesn’t exist or isn’t accurate for many fish species.

Age validation and data generation for fisheries stock assessments are routine^10–12^, with most data estimated by sectioning fish ear stones (otoliths) and counting growth rings (i.e., annuli) associated with temperature-dependent seasonal growth rates. Otolith ageing is time- and labor-intensive, necessitates sacrificing individual fish to extract otoliths, and is only applicable to bony fishes (i.e., not sharks or other chondrichthyans). Additionally, some species have small or difficult-to-extract otoliths, which has led researchers to use alternative calcifying structures for age determination^13^, but these different methodologies can create inconsistencies in ageing assessments within a single species^14,15^. Long-lived species that show a dramatic decrease in growth rate later in life have compressed annuli that are difficult to distinguish, leading to inaccurate age data^8^. In some cases, commercially important fishes (e.g., goosefishes, *Lophius* spp., Lophiidae) do not produce consistent annuli on any calcifying structure^16^. Because of these nuances associated with otolith ageing, there is a need for a more universally applicable method to accurately assess age in fish populations. Age-related epigenetic signatures have emerged as one of the most accurate biomarkers of age across a wide range of animal species, presenting an opportunity to develop an epigenetic ageing tool for fish as an attractive alternative to lethally sampling calcifying structures. ‘Epigenetics’ refers to processes that alter gene regulation without changing the DNA sequence itself^17^. DNA methylation is a key epigenetic mechanism involving the addition of a methyl group to a cytosine resulting in the formation of 5-methylcytosine, which predominantly occurs in DNA sequences where cytosine is immediately followed by guanine (CpG sites). Changes in DNA methylation patterns can occur in response to the environment^18–20^, but there are portions of the genome that consistently change in methylation level with chronological age, a conserved mechanism across a variety of taxa^21,22^. Exploration of age-associated DNA methylation patterns at CpG sites has resulted in species-specific ageing clocks for humans^23^, elasmobranchs^24^, marine mammals^25^, and lobsters^26^ as well as ‘universal’ ageing clocks, like for 128 mammalian species^27^ and a microarray platform across various mammalian species (‘HorvathMammalMethyl-Chip40’^28^). Over the last five years, epigenetic ageing clocks have been published for ten species of teleost fish, including Golden perch (*Macquaria ambigua*)^29^, Zebrafish (*Danio rerio*)^30^, Atlantic halibut (*Hippoglossus hippoglossus*)^31^, Japanese medaka (*Oryzias latipes*)^32^, European seabass (*Dicentrarchus labrax*)^33^, Red grouper (*Epinephelus morio*)^34^, Northern red snapper (*Lutjanus campechanus*)^34^, Blackbelly rosefish (*Helicolenus dactylopterus*)^35^, Mangrove rivulus (*Kryptolebias marmoratus*)^36^, and Turquoise killifish (*Nothobranchius furzeri*)^37^. These clocks represent eight teleost fish orders and were generated from six different tissue types. Nonetheless, no epigenetic clocks exist for a species within the Family Gadidae, a particularly important group of groundfishes in the Northern Atlantic Ocean.

Haddock (*Melanogrammus aeglefinus*, Gadidae) is a commercially important fishery in the North Atlantic, producing 300,000 tons of fish annually, with landings reported by 23 different nations over the last decade^38^. In the United States, haddock supports active commercial and recreational fisheries across the Gulf of Maine and Georges Bank, where landings total ∼7,000-10,000 tons each year. Fisheries-independent monitoring for this species in the United States includes bottom trawl surveys that sample the majority of the 12-year lifespan (up to ∼10 years old). Recent data from these surveys have shown a greater than normal proportion of older individuals due to an unprecedented high recruitment event in 2013^39^. This unique age structure allows for a higher number of samples from older individuals, often a limiting factor in epigenetic ageing clock generation^40^. Haddock satisfies additional resource requirements for building an epigenetic ageing clock, like a reference genome^41^ and high confidence in accurate otolith ages for clock validation^8,42^. This study took advantage of the timely availability of older haddock individuals to create a species-specific epigenetic ageing clock. The haddock clock generated here is only the second fish epigenetic ageing clock developed from whole genome sequencing (along with Atlantic halibut^31^), offering deeper insight into the genomic distribution and functional significance of age-associated DNA methylation patterns.

## Results

### Haddock otolith ageing and demographics

Haddock individuals (n= 277) collected in the Gulf of Maine and Georges Bank during NOAA’s Fall 2022 and Spring 2023 trawl surveys ranged from 16 cm – 68 cm in length (Figure 1a-b) and 0.040 - 3.155 kg in weight (Supplemental Figure 1; Supplemental Table S1). Decimal age of individuals, determined by otolith annuli counting, ranged from 0.7-10.2 years with rapid growth over the first 5 years before growth rate stabilizes in older individuals. There was a wide range of size classes at each age interval after one year of age, demonstrating the challenges of using length as a proxy for age. There was no sexual dimorphism seen in younger individuals, however, female older individuals displayed increased length and weight values at the end of their lifespan compared to males (Supplemental Table S2).

**Figure 1.**
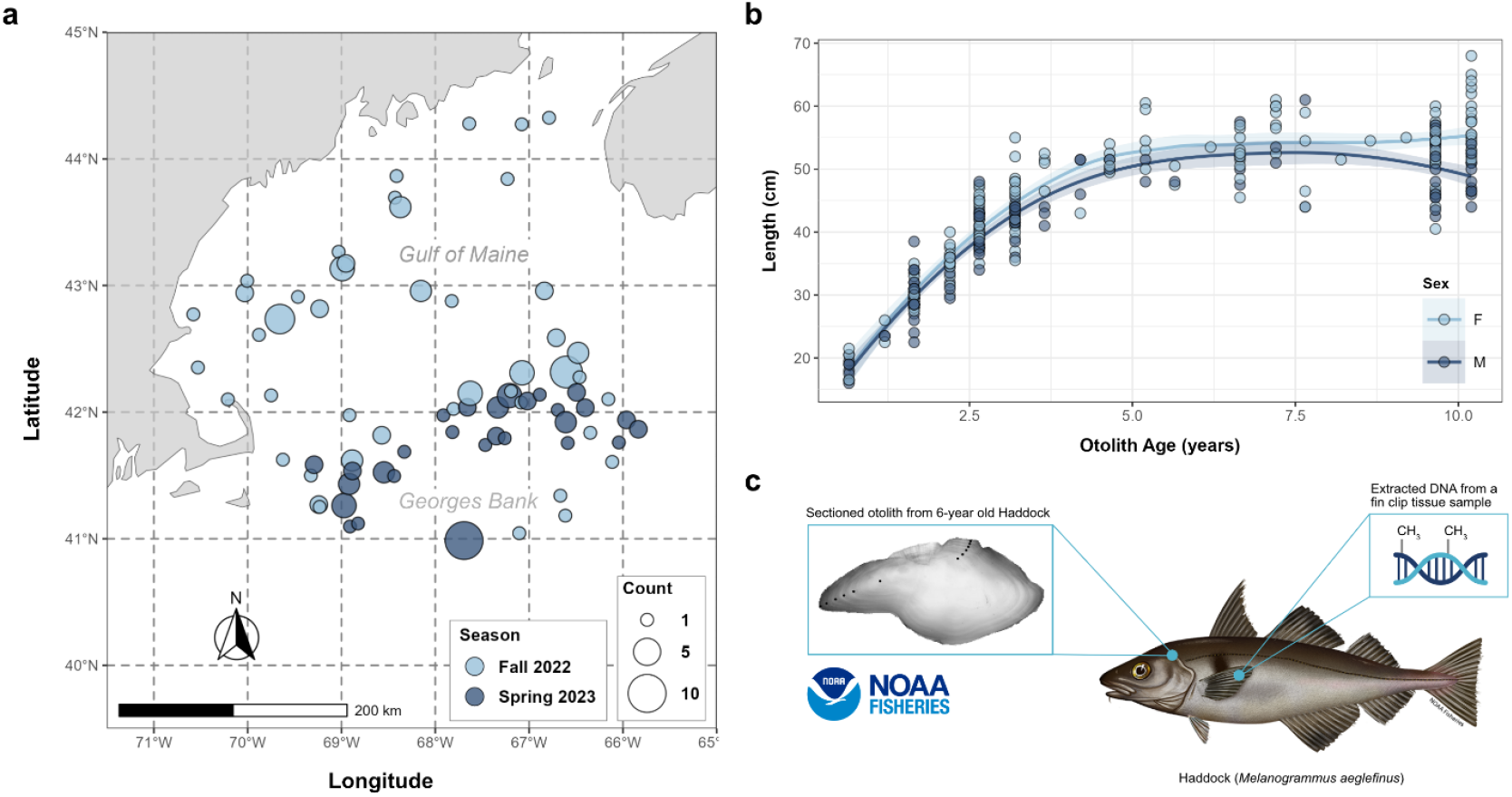
Sampling scheme for haddock biological collection. a) Locations of each collection site for Fall 2022 (light blue) and Spring 2023 (dark blue) Bottom Trawl surveys. Size indicates the number of fish collected from each site. b) Otolith age (years) compared to fish total length (cm) from collected individuals (n=277) during both trawl surveys. c) Graphic depiction of extracted DNA and methylation patterns from haddock along with a sectioned otolith from a 6-year-old individual with annuli marked by black circles.

### Sequencing quality, bisulfite conversion, and global methylation levels

Sequencing data for 140 individuals selected to represent the entire age range with an equal number of males and females in each age category, produced high quality data (>Q30) with an average of 152.3±2.35 million reads (Supplemental Table S3-4). On average, 44.13±0.39% of the reads aligned to the scaffold-level reference genome and 64.73±0.11% of cytosines in a CpG context were methylated (Supplemental Figure 2). Median bisulfite conversion efficiency across both sequencing runs was 97.72±0.89%, and 133 individuals (n=91 training; n=42 testing) with >96% conversion efficiency were used in methylation analysis (Supplemental Figure 3; Supplemental Table S5-6). As a result, samples contained 1,139.19±26.65 million cytosines for subsequent analysis. Global CpG DNA methylation increased slightly, but significantly (p < 0.0001), with age during the first five years and then remained constant through age 10.2 years (Supplemental Figure 4). This pattern was more pronounced in male individuals, corresponding to similarly timed increase in length (Figure 1) and weight (Supplemental Figure 1). Total methylation (CpG + CHG + CHH) level was 76.06±0.26% across all samples and the median CpG loci methylation level was 75.67±0.003%. These values are consistent with those previously reported for teleost fish, which exhibit ∼60-80% of their genome methylated globally^43^ and CpG loci methylation levels >50%^43^.

### Generation and validation of an epigenetic ageing clock for haddock

Filtering CpG sites to those present in ≥90% of samples yielded 813,743 loci and those present in 100% of samples yielded 108,411 loci per sample. After removing SNPs, low variance, and low confidence loci (i.e., <10% range across samples and all samples <10% level), the ≥90% presence set contained 642,346 loci and the 100% presence set contained 105,602 loci. The Bayesian GLMM identified 47,373 loci from the ≥90% presence and 7,731 loci from the 100% presence sets that were significantly associated with age. Both sets of loci were tested within the lasso and elastic net model workflow to assess the most accurate epigenetic ageing clock. The multi-iteration lasso model further reduced this set to 1,263 loci to be used in the final elastic net regression model function. The most effective epigenetic ageing model included 111 CpG loci with an R^2^=0.97 and mean absolute error (MAE) of 0.12 years (Figure 2a). The median MAE for the training set was 0.09 years and 0.68 years for the testing set of individuals (Figure 2b). Overall, this model predicted age within 1 year at 93% accuracy (i.e., 7 individual predictions were 1+ year away from validated age). The final epigenetic ageing clock contained CpG loci with DNA methylation patterns that either increased (n=66) or decreased (n=45) with age (Figure 2c). Although the final ageing model reduced the number of features needed to predict age to 111, this workflow was repeated many times during this analysis, which produced thousands of models. A ‘core’ set of 3,797 CpG loci was identified as sites that were consistently included in the top ranked models. The best ageing clock is presented in Figure 2, but there were many iterations that accurately predicted age (e.g., R^2^>0.95) using a combination of the ‘core’ set (Supplemental Table S7-9).

**Figure 2.**
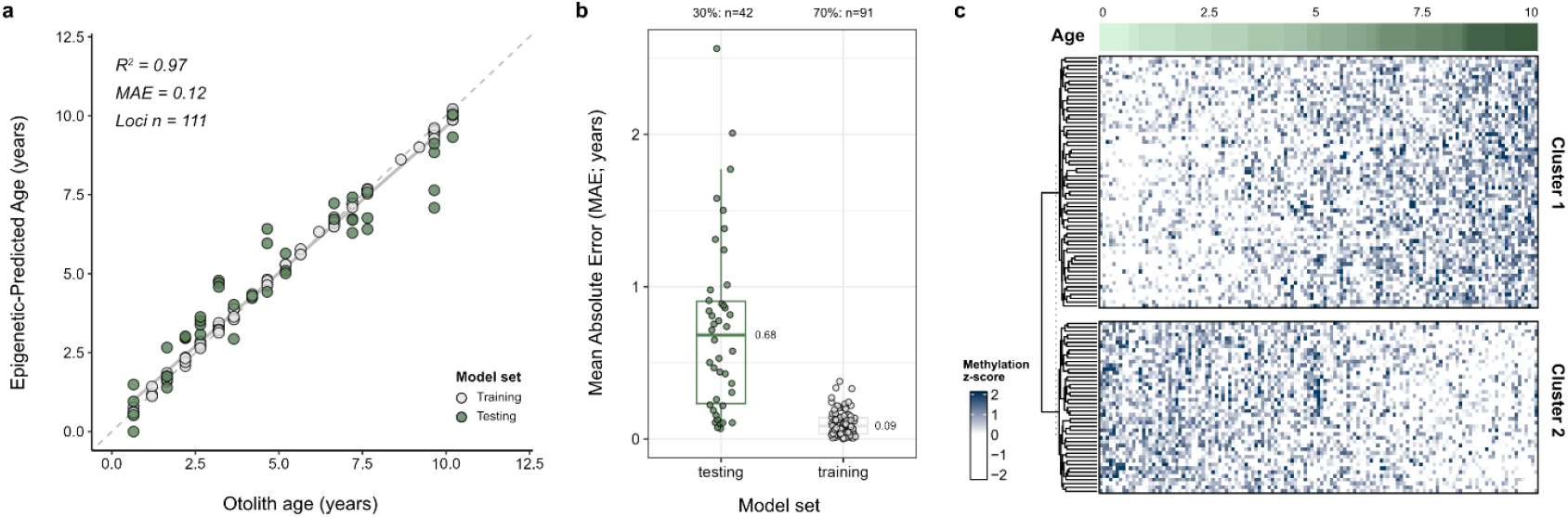
A species-specific haddock epigenetic ageing clock generated from 111 CpG loci. a) Otolith age (years) compared to epigenetic-predicted age (years). Ageing model yielded R^2^=0.97 with a mean absolute error (MAE) of 0.12 years. b) MAE for all samples from 70% training (n=91 fish) and 30% testing (n=42 fish) sets. Median training MAE calculated as 0.09 years and testing MAE as 0.68 years. c) Scaled DNA methylation levels (z-score) for the 111 CpG loci (rows) for all 133 individuals (columns). Cluster 1 represents loci that increase methylation levels with age and Cluster 2 represents loci that decrease methylation levels with age.

### Genomic location of age-related sites

CpG loci significantly correlated with age occurred throughout the genome in non-genic and genic regions (Figure 3a). Out of the 47,373 CpG loci in the ≥90% presence set, 859 appeared in 3’ untranslated region (UTR), 840 in 5’ UTR, 10,254 in CDS, and 21,750 in intron genic regions as well as 608 in promoter, 609 in terminator, and the remaining 12,453 in other non-genic regions. Of the 111 CpG loci included in the final ageing clock, 3 were in 3’ UTR, 2 in 5’UTR, 31 in CDS, 45 in introns, 1 in promoter, and 29 in other non-genic regions (Supplemental Table S10). Genic regions held the majority of age-related CpG loci in the final ageing clock (72.97%) and the larger age-associated set (71.14%). The ≥90% presence set of CpG loci were found across 5,518 of the 8,420 scaffolds in the haddock genome. The number of loci per scaffold varied from 1-186 with larger scaffolds generally containing a greater number of CpGs (Figure 3b).

**Figure 3.**
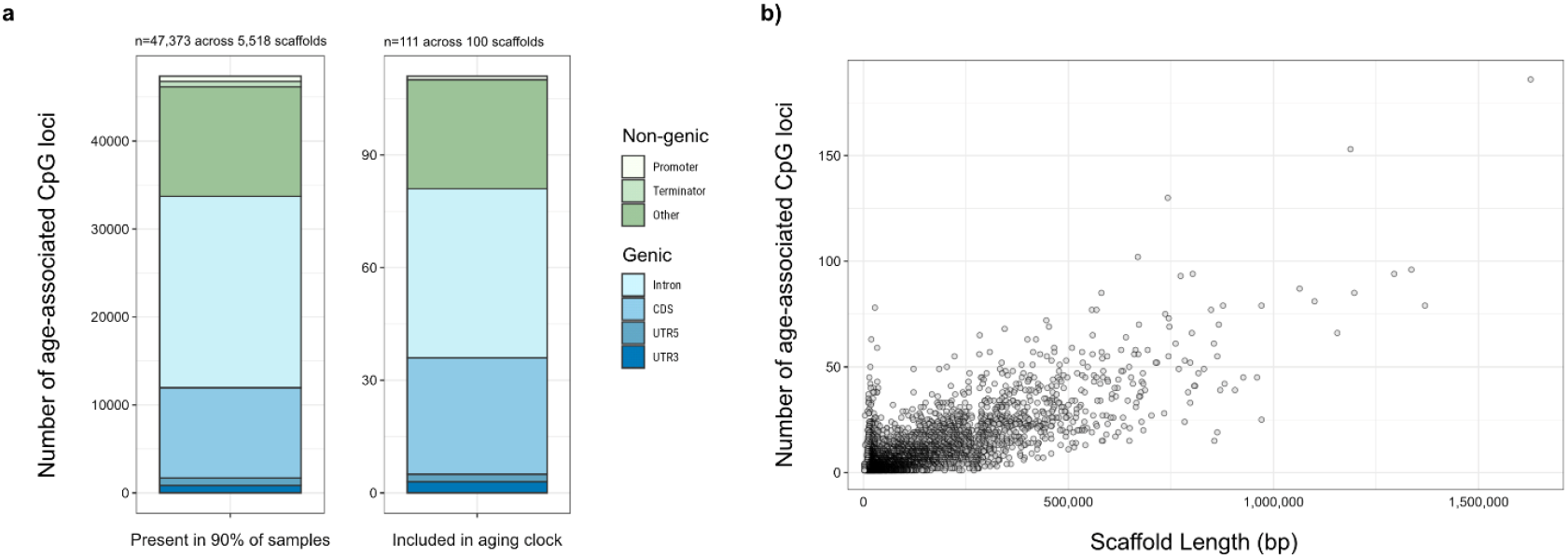
Region information and genomic location of the significantly age-associated CpG loci. A) The number of CpG loci from the ≥90% presence set (n=47,373 loci from 5,518 different scaffolds) and from the final ageing clock (n=111 loci from 100 different scaffolds) located in non-genic (e.g., promoter, terminator, other) and genic (e.g., intron, CDS, 5’ UTR, 3’ UTR) regions. B) The number of CpG loci within each of the 5,518 scaffolds from the ≥90% presence set.

### Functional annotation of genes containing age-associated CpG loci in haddock

Age-associated CpG loci from the ≥90% presence set that increase in methylation levels (%) with age (‘Cluster 1’; n=28,762 loci) and decrease in methylation level (%) with age (‘Cluster 2’; n=18,611 loci) were subsetted to those within genic regions. ‘Cluster 1’ contained 14,464 genes with 6,785 protein sequences, and ‘Cluster 2’ contained 9,881 genes with 5,012 protein sequences. Genes were further filtered to those with >10 CpG loci to focus on the most age-related functions, which resulted in 395 genes with 291 protein sequences. These protein sequences were analyzed in STRING to assess protein-protein interaction (PPI) networks and conduct functional enrichment analysis. The resulting network was comprised of 240 nodes with 55 edges (PPI enrichment p-value < 0.0001) and 11 sub-networks. The largest sub-network contained 19 nodes and 30 edges (PPI p-value < 0.0001), with 195 significantly enriched terms from Reactome pathways, Gene Ontology, KEGG pathways, and protein domains (Supplemental information; Figure 4a-b).

**Figure 4.**
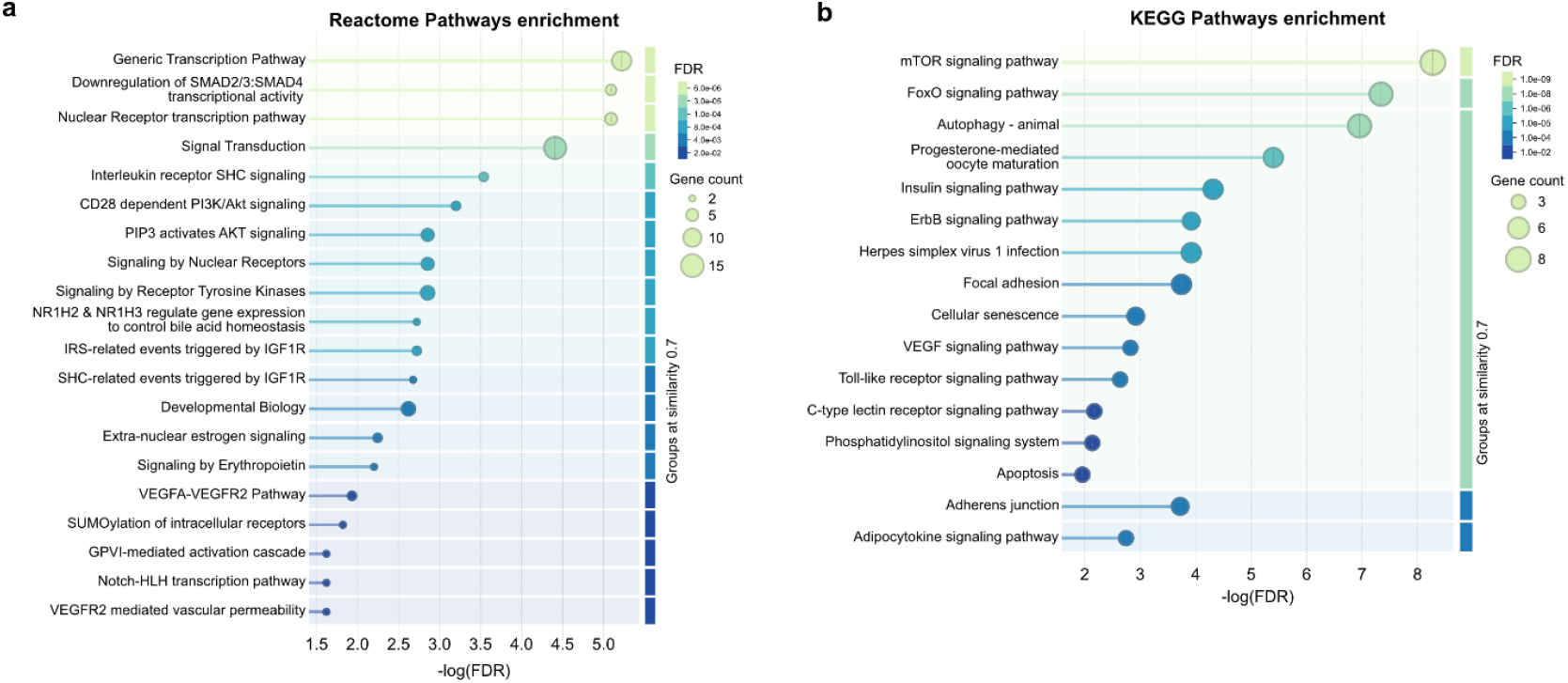
Significantly enriched functions from genes with >10 age-related CpG loci. Each point is a set of genes (size indicative of number of genes) associated with a significantly enriched term. Gene sets are grouped at 80% similarity with false discovery rate (FDR; i.e. p-values corrected via Benjamini-Hochberg procedure) shown by color and ranked by signal (i.e., weighted harmonic mean between the observed/expected ratio and -log(FDR). Functions included in a) Reactome pathways. The 45 Reactome pathway terms were merged by similarity = 1.0 resulting in the 20 terms shown here. b) KEGG pathways.

Within Reactome pathways, functions related to metabolism like phosphatidylinositol (PI) metabolism (MAP-1483255) and synthesis of PIPs at the plasma membrane (MAP-1660499) along with developmental pathways like adipogenesis (transcriptional regulation of white adipocyte differentiation; MAP-381340) and nervous system development (RET signaling; MAP-8853659) were significantly enriched. However, most of the significantly enriched functions were related to the immune system, signal transduction, and gene expression and transcription (Figure 4a; Supplemental information). In the immune system, genes with age-related CpG loci were found in interleukin signaling pathways (Interleukin-2 family [MAP-451927]; Interleukin-3, Interleukin-5, and GM-CSF [MAP-512988]; Interleukin receptor SHC [MAP-912526]; Regulation of signaling by CBL [MAP-912631]) and adaptive immune system functions related to the regulation of T cell activation by the CD28 family (Co-stimulation by CD28 [MAP-389356, MAP-388841]; CD28 dependent PI3K/Akt signaling [MAP-389357]). There were five major signal transduction pathways (MAP-162582) represented: signaling by erythropoietin, receptor tyrosine kinases, nuclear receptors, transforming growth factor beta (TGFB) family members, and intracellular second messengers. Signaling by erythropoietin (MAP-9006335) was specific to erythropoietin activating phosphoinositide-3-kinase (PI3K; MAP-9027276). Receptor tyrosine kinases (MAP-9006934) were dominated by two groups of growth factors: vascular endothelial growth factors (VEGF [MAP-194138]; VEGFA-VEGFR2 Pathway [MAP-4420097]; VEGFR2 mediated vascular permeability [MAP-5218920]) and type 1 insulin-like growth factor 1 receptors (IGF1R [MAP-2404192]; IGF1R signaling cascade [MAP-2428924]; SHC-related events triggered by IGF1R [MAP-2428933]; IRS-related events triggered by IGF1R [MAP-2428928]). Nuclear receptor signaling included Extra-nuclear estrogen signaling (MAP-9009391) and NR1H2 and NR1H3-mediated signaling (MAP-9024446) like NR1H3 & NR1H2 regulate gene expression linked to cholesterol transport and efflux (MAP-9029569) and NR1H2 & NR1H3 regulate gene expression to control bile acid homeostasis (MAP-9623433). TGFB family member signaling (MAP-9006936; TGF-beta receptor complex [MAP-170834]) was specifically related to SMAD transcriptional activity (SMAD2/3:4 heterotrimer [MAP-2173793]; Downregulation of SMAD2/3:SMAD4 [MAP-2173795]). Finally, intracellularly second messenger (MAP-9006925) signaling was dominated by PIP3 activating AKT signaling (MAP-1257604) functions like negative regulation of the PI3K/AKT network (MAP-199418) and PI5P, PP2A and IER3 Regulate PI3K/AKT Signaling (MAP-6811558). Apart from signaling transduction, age-related CpG loci were involved in genes related to gene expression and transcription regulation (MAP-74160; MAP-212436) and several pathways involving RNA Polymerase II transcription (MAP-73857) like nuclear receptor transcription (MAP-383280), notch-HLH transcription pathway (MAP-350054), and regulation of TP53 expression and degradation (MAP-6806003; MAP-6804757) (Supplemental Table S11-12).

Many of the signaling functions described above are involved in enriched KEGG pathways like mammalian target of rapamycin signaling (mTOR; map04150), Forkhead box O signaling (FoxO; map04068), insulin signaling (map04910), Erythroblastic oncogene B signaling (ErbB; map04012), PI signaling (map04070), and VEGF signaling (map04370) (Figure 4b). Additional enriched KEGG pathways included immunity-related functions like toll-like receptor signaling (map04620) and C-type lectin receptor signaling (map04625). Overall, these enriched Reactome and KEGG functions were involved in processes like autophagy (map04140), progesterone-mediated oocyte maturation (map04914), focal adhesion (map04510), adherens junction (map04520), infection signaling (Herpes simplex virus 1 infection; map05168), cellular senescence (map04218), regulation of actin cytoskeleton (map04810), and apoptosis (map04210) (Supplemental Table S11-12).

## Discussion

This study presents a highly accurate, species-specific epigenetic ageing clock for haddock that is independent of sex or length. This is the first published epigenetic ageing model for a representative of the Family Gadidae, a commercially important teleost fish family worldwide^38^ and is the second to generate a fish ageing model from whole genome sequencing^31^ reflecting an opportunity to investigate functional implications of genome-wide epigenetic ageing signatures.

Although debated^44^, global DNA methylation patterns tend to decrease with age^45,46^, which has been demonstrated in fish, including killifish (*Nothobranchius furzeri*)^47^ and grass carp (*Ctenopharyngodon idellus*)^48^. In contrast, this study found that whole genome CpG global methylation increased from ages 0 to 5 and then stabilized, closely mirroring growth dynamics. The rapid methylation increase during young year-classes was present in both sexes, but much more pronounced in males relative to females. Haddock do not generally exhibit sexual dimorphism regarding pectoral fin growth or overall length^49^, and no sex-based differences in length or weight were observed in our data during the rapid methylation increase observed in the first half of life. Despite the lack of phenotypic differences between males and females, this study documented global DNA methylation pattern differences among sexes during juvenile development. In fish, development and sex determination can be heavily epigenetically regulated^50–52^, specifically with respect to growth-related genes that show distinct sexual dimorphism with minimal similarity in DNA methylation patterns between males and females^53–55^. Thus, there are likely differential energetic, and therefore epigenetic, needs associated with sexual dimorphism that could lead to the sharper increase in global methylation seen in males.

Unlike global DNA methylation patterns, site and region-specific CpG methylation changes with age are known to be a mix of hyper-methylation^32,37,56^ and hypo-methylation^34,37,57,58^. Aligning with previous work, the age-related CpG loci found in haddock in this study included both hypermethylated and hypomethylated directional change. Regardless of hypo- or hyper-methylation, DNA methylation acts as a regulatory mechanism for gene expression, which can have large functional and phenotypic consequences in fish^59–61^. This study found that as fish age, methylation levels on CpG loci significantly changes within genes related to transcriptional regulation along with innate and adaptive immune system, signal transduction, metabolism, and development. DNA methylation pattern changes within the fin tissue may be needed to more effectively regulate transcription with age, particularly in epithelial cells that are the first line of defense against foreign molecules, infections, and environmental stimuli. Along with changes to external signal regulation, older fish cell membranes exhibit two major physical changes related to membrane integrity: increase in cellular deformities^62^ and dysregulation of lipid metabolism, which is a major cause of age-related diseases^63^. Other studies have found epigenetic reprogramming to be a mechanism underlying the cell’s response to these disruptions in lipid metabolism^64^ and cell membrane integrity^62,65^. Ultimately, DNA methylation may play an important role in maintaining cellular function and the ability to respond to their environment as fish age.

By taking advantage of these epigenetic changes within the fin tissue, this study generated an epigenetic clock model from a sample set representative of a broad geographic area across the Northwest Atlantic. The model was subsequently validated on 40+ individuals from a separate testing sample set (30% of total samples). The clock demonstrated strong performance, predicting age ±1 year with 93% accuracy for all samples, suggesting that this tool will be reliable for age estimation of haddock individuals across the Northwest Atlantic. The final clock uses 111 CpG loci from across the genome, consistent with current epigenetic ageing clocks for marine organisms which use ∼20 – 200 CpG loci^32,34,66,67^. This tool can now be used by the community for ageing haddock by doing targeted bisulfite sequencing of the 111 CpG loci and processing percent methylation per CpG data with the analysis pipeline we developed. Besides applications for haddock, this larger dataset of age-related CpG loci (n=47,373) derived from whole genome sequencing can serve as a platform for epigenetic ageing clock development in other species, particularly in closely related, commercially important species like Atlantic Cod (*Gadus morhua*), Pollock (*Pollachius* sp.), and Whiting (*Merlangius merlangus*).

This method is a powerful tool for fisheries management to generate age information from fin tissue in a cost-effective, scalable, and non-lethal way. As epigenetic ageing model generation gains popularity, the applicability and consistent accuracy of the produced clock will be increasingly important. Implementation of epigenetic ageing methods into fisheries management practices will require clocks to be reproducible, accurate through time, and robust against environmental influences. We recommend that the most reproducible model is one that is 1) built using a large sample set that is representative of the population to which the tool will be applied, and 2) evaluated on samples not used to build the model (i.e., testing set, commonly 30%). Recent evidence has shown that when an epigenetic clock is developed using tissue samples from a specific population, and subsequently applied to a different population, the accuracy of chronologic age estimation decreases^68^. The machine learning methods used in epigenetic ageing clocks are prone to overfitting training data and thus increases the risk that a model is not usable for a broader population. Epigenetic ageing methods in humans and other mammals have shown this exact predicament – several clocks will yield different results for the same species^69^. Because DNA methylation patterns are responsive to the environment, including factors like temperature, geographic region, and captivity^70–72^, the sample set and CpG loci used in clock generation are both extremely important to ensuring the model will withstand environmental pressures and prove useful many years later. Several recent studies found their ageing clocks were not impacted by environmental stressors^32,73^, which provides some confidence that dynamic environments may not disrupt the tool’s utility.

In the last five years, the field has established the utility and effectiveness of piscine epigenetic ageing clocks across the fish tree of life – from golden perch (*Macquaria ambigua*)^29^ to turquoise killifsh *(Nothobranchius furzeri*)^37^. To date, only a handful of marine-focused studies have made strides towards multi-tissue^35,74^ and multi-species clocks^74–77^. Most of these studies were generated for marine mammals, leaving piscine multi-species clocks yet to be established. Given the large evolutionary history within teleost fish^78^, the feasibility of a ‘universal’ piscine clock remains unknown. Thus far, epigenetic ageing clocks have been limited to species with highly reliable traditional ageing methods for validation purposes. However, epigenetic ageing approaches may act in concert with other modern ageing methods like fourier-transform near-infrared spectroscopy (FT-NIRS), which correlates spectral signatures with age^79^. Although lethal, FT-NIRS may provide an increased sample size potential that is cost-effective and reduces labor requirements. Vice versa, validated epigenetic ageing clocks may serve as an alternate, non-lethal, and cost-effective validation method while building FT-NIRS protocols. Along with advancing methods technology, the largest impact of this tool will be for species, like goosefish (*Lophius americanus*, Lophiidae) that lack any method for reliably estimating age^10^ or gadoid fisheries in brackish waters like the Baltic Sea that lack the visual contrast within otolith growth rings because of the low salinity^80^. Creating a widely applicable, ‘universal’ piscine clock would vastly improve models used in sustainable fisheries management efforts.

## Methods

### Sample collection

Fin clips and otoliths were removed from 277 haddock individuals collected during the Fall 2022 and Spring 2023 in the Georges Bank and the Gulf of Maine regions (Figure 1a), as part of a regular bottom trawl survey operated by National Oceanic and Atmospheric Administration’s Northeast Fisheries Science Center (NEFSC)^81^. During sampling, total length (0.5 cm), weight (0.001 kg), and sex (M/F) were measured, and the pectoral fin was removed with sterilized clippers, preserved in 100% molecular grade ethanol, and immediately stored at -20°C on the boat. Individual fish fin clips were brought to Gloucester Marine Genomics Institute (GMGI) and stored at -80°C prior to processing. Otoliths were removed and stored dried in an envelope prior to age analysis at NEFSC.

### Otolith age prediction

Haddock were aged using sectioned otoliths. Otoliths were cut twice using a high-speed saw with a single metal-bonded diamond blade that resulted in a 0.18 mm thick section encased in resin. Each otolith was viewed under a binocular microscope using reflected light at 15-25X magnification. Age was calculated by counting the number of hyaline zones from the nucleus to the edge of the otolith^82^.

### Sample selection

From the total sample set (n=277), 140 individuals representative of all ages were selected for DNA extraction and sequencing. Age bins were established assuming a born-on date of March 1^st 83^ and generating a decimal age based on collection date. Given that collection dates were not protracted within a season (23 days in the fall and 11 days in spring), spring fish were assigned a decimal age of otolith age + 0.2 (mean=0.21±0.009) and all fish collected in the fall were assigned a decimal age of otolith age + 0.7 (mean=0.69±0.015), which resulted in half-year age bins from 0.7 to 10.2 years. Within each age bin, individuals were chosen across a stratified length range with the subset including ∼50% males and ∼50% females to ensure a representative set. This sample scheme was intentionally chosen to limit bias for a particular age class, sex, or length while ensuring sufficient sample size to construct an epigenetic clock^40^.

### DNA Extractions

Fin clips were wiped down with sterile kimwipes to clean off any debris or mucus and a ∼3×3 mm piece (∼20 mg) was cut with clippers sterilized in 10% bleach, 70% ethanol, and MilliQ water to be used for DNA extraction. This small tissue clipping was used in the Qiagen DNeasy Blood & Tissue Kit DNA extraction kit (Qiagen, Cat. #69506) according to manufacturer’s instructions with the following additions. 20 µL of 10 mg/mL RNase A was added to each sample after lysis and incubated for 3 minutes at room temperature prior to the addition of Buffer AL. Two elution steps were performed by adding 50 µL of AE buffer to each spin column with a 3-minute incubation prior to centrifuging. DNA quality was assessed using gel electrophoresis (1% agarose gel, 45 minutes, 2A, 100V) and quantity (ng µL^-1^) was measured using an Invitrogen Qubit 4 Fluorometer (ThermoFisher, Cat. #Q33238) with a dsDNA Broad Range assay kit (ThermoFisher, Cat. #Q32853).

### Whole genome bisulfite sequencing

Extracted DNA (20 ng) was bisulfite-converted and prepared for Whole Genome Bisulfite Sequencing (WGBS) using the Pico Methyl-Seq Library Preparation Kit (Zymo Research, Cat. #D5456) according to manufacturer’s instructions with the following modifications. Each sample included a 2.5% (0.5 ng) spike-in of *E. coli* non-methylated genomic DNA (Zymo Research, Cat. #D5016) to later assess bisulfite conversion efficiency. After the second addition of M-Wash Buffer in column clean-up steps, samples were centrifuged for 2 minutes at 16,000 rcf. DNA elution buffer was warmed at 56°C prior to use and once added to the spin columns. Samples were incubated at room temperature for 5 minutes before centrifugation to increase DNA yield. Following bisulfite conversion, the first amplification step included 8 PCR cycles, and the second amplification included 10 PCR cycles. Unique Dual Indices (UDI) were added to each sample using 2 µL of combined i5 and i7 index primers (Zymo Research, Cat. #D3096) during the second amplification. Resulting PCR product was cleaned with equal volume (25 µL) of PCR Clean DX beads (Aline Biosciences, Cat. #C-1003) and 80% ethanol. Cleaned samples were resuspended in 15 µL of room temperature DNA elution buffer. Libraries were quantified using an Invitrogen Qubit 4 Fluorometer (ThermoFisher, Cat. #Q33238) with a dsDNA High Sensitivity assay kit (ThermoFisher, Cat. #Q32854). Quality was assessed using an Advanced Analytical Fragment Analyzer with a NGS High Sensitivity Kit (Agilent, Cat. #5191-6578). Libraries that showed a single, smooth peak in DNA fragment size ranging from 100-500 bp were pooled in equimolar amounts. All samples were sequenced in two batches across four flow cell lanes for 2 × 150 bp sequencing on an Illumina NovaSeq S4 (Illumina NovaSeq 6000 S4 Reagent Kit v1.5 300 cycles, Cat # 20028312) at the University of Connecticut’s Center for Genome Innovation.

### WGBS bioinformatic workflow

Sequencing files were analyzed on Northeastern University’s High Performance Discovery Cluster. Quality of raw files were initially assessed using FastQC (v0.11.9) and MultiQC (v1.16)^84^ to determine the best trimming parameters. Based on the MultiQC report, different 3’ and 5’ end clipping parameters were tested within the following methylation calling pipeline to optimize processed read quality and reduce m-bias. TrimGalore! (https://github.com/FelixKrueger/TrimGalore; v0.6.5) was used to trim paired-end reads and remove Illumina adapters with the following parameters: `--clip_r1 10`, `--clip_r2 10`, `--three_prime_clip_r1 10`, and `--three_prime_clip_r2 10`. Trimmed reads were then aligned to the haddock genome^85^ using Samtools (v1.9)^86^ and the Bismark (v0.24.2)^87^ align function with the following parameters: `--relax_mismatches`, `-- num_mismatches 0.6`, and `--non-directional`. Aligned bam files were deduplicated with the Bismark `deduplicate_bismark` function and the methylation calls were extracted with the Bismark `bismark_methylation_extractor` function. The top and bottom strand were merged with the Bismark `coverage2cytosine` function with the following parameters: `--merge_CpG` and `-- zero_based`. Reads were further filtered to CpG sites found in ≥90% of samples with 10X coverage using `intersectbed` and `multiIntersectBed` functions from bedtools (v2.29.0)^88^. The resulting bed files were analyzed with R and RStudio (v4.2.0) on the Discovery Cluster. Global DNA methylation level (the percentage of cytosines methylated across the genome) was calculated within the Bismark methylation extraction step.

### Calculating bisulfite conversion efficiency

Trimmed reads were aligned to the *Escherichia coli str. K_12 substr. MG1655* (Genome assembly ASM584v2; RefSeq assembly accession: GCF_000005845.2) using the same steps and parameters as described above. The bisulfite conversion efficiency was calculated for each library by dividing the sum of unmethylated cytosines in the CHH and CHG context by the total cytosine in the CHH and CHG context that were analyzed: (U CHH + U CHG) / ((U+M CHH) + (U+M CHG)). Samples with <96% conversion efficiency were not included in the ageing model generation.

### CpG loci filtering

DNA methylation level (%) was calculated for each CpG locus by dividing the number of methylated reads by the total number of reads. Loci with low variance were filtered out using the ‘nearZerovar’ function from the R package caret (v6.094)^89^ with a frequency cut-off ratio of 85/15 (`freqCut = 85/15`) and a maximum percentage of unique values of 50 (`uniqueCut = 50`). Single nucleotide polymorphisms (SNPs) were identified from the merged bam files with BS-Snper (v1.0)^90^ with the following parameters prior to removal: `--minhetfreq 0.1, --minhomfreq 0.85, -- minquali 15, --mincover 10, --maxcover 1000, --minread2 2, --errorate 0.02`, and `--mapvalue 20`. CpG loci were additionally filtered out if DNA methylation levels were under 10% or if the range of percent methylation was less than 10% across all samples.

### Epigenetic age prediction

The resulting DNA methylation data matrix was then used for downstream age prediction analyses. To identify genomic sites with age-correlated DNA methylation, a Bayesian approach was used to estimate parameters of a generalized linear mixed model (GLMM) with decimal otolith age as a fixed factor and sex as a random factor. The `stan_glmer` function from the rstanarm package (v2.26.1)^91^ was used with the following model: `stan_glmer(matrix(c(M, U), ncol=2) ∼ Age + (1 | Sex)`; where M represents the number of methylated reads and U represents the number of unmethylated reads at each site. The model was run with 8,000 iterations (iter=8000) and family set to binomial (family = binomial; link = logit). CpG sites with models that converged (Gelman-Rubin ‘rhat’ statistic < 1.010) or had an effective size of >2,000 were removed. Of the remaining models, any with a p-value <0.05 were considered significantly correlated with age. This analysis was performed on CpG sites present in ≥90% of samples and 100% of samples. Imputations of missing data were conducted using the Multivariate Imputation by Chained Equations R package (mice; v3.16.0)^92^. The top 20% CpG sites most significantly related to age from the ≥90% presence set were used to predict epigenetic age in subsequent models.

To evaluate the prediction capabilities of the model, the samples were first divided into ‘training’ and ‘test’ sets by randomly assigning 70% of samples to a ‘training set’ and the remaining 30% to a ‘test set’ using 100 iterations of a bootstrap function. Several iterations of lasso and elastic models were employed to identify the optimal number of CpG sites predictive of age. An iterative analysis was used for each training set to repetitively retain only the top 75% most predictive age-correlated CpG sites using 7 iterations of a lasso regression model. This was performed with the ‘cv.glmnet’ function from the glmnet package (v4.1-8)^93^ with the following parameters: alpha=0, nfold=10, family= “gaussian”, type.measure= “mae”, and standardize=false. The resulting CpG loci were used as input for a function that randomly subsets the input to n=650 over 500 iterations, followed by running ‘cv.glmnet’ as described above with alpha=0.02, and then predicting the age of the testing samples. The function selects the top model (out of 500) with the lowest mean absolute error (MAE) calculated from the testing set of samples. Throughout this analysis, alpha values of 0, 0.01, 0.02, 0.1, and 0.5 were tested to evaluate model performance and number of sites retained. Linear regressions were used to compare the otolith age to epigenetic-predicted age. The top models were those with the lowest value of mean absolute error (MAE) and highest R^2^ value.

### Genomic location and function

The CpG loci significantly associated with age (i.e., output from the Bayesian general linear mixed model) from both the ≥90% and 100% presence datasets, and more specifically, the loci included in the final epigenetic ageing clock were analyzed to determine genomic location and function. These CpG loci lists were compared to the genome gff file^85^ using the GenomicRanges (v.1.56.2)^94^ and IRanges (v.2.38.1)^94^ R packages to identify non-genic, promoter (i.e., 200 bp upstream of gene start), terminator (i.e., 200 bp downstream of gene end), coding sequence (CDS), exon, 3’ untranslated region (UTR), 5’ UTR, and intron regions. DNA methylation directional change (i.e., increase or decrease with age) was assessed by calculating z-score for each CpG loci and applying a k-means calculation to identify two clusters. The CDS regions from genes with >10 age-associated CpG loci were extracted from the genome protein fasta file^85^ and analyzed in STRING (v12.0)^95^. With no haddock reference available on STRING, Atlantic Cod, *Gadus morhua*, was used as a reference and sub-networks of interacting genes with a minimum interaction score of 0.7 (high confidence) were identified using k-means clustering. STRING identifies significantly enriched gene functions (e.g., gene ontology), pathways (e.g., Reactome and KEGG), and protein families (e.g., UniProt, Pfam, SMART, and InterPro) within each network.

## Acknowledgements

We would like to thank the National Oceanic and Atmospheric Administration (NOAA) Northeast Fisheries Science Center (NEFSC) for collecting and providing these samples from the Fall 2022 and Spring 2023 Bottom Trawl Surveys.

## Author Contributions

ELS conducted the epigenetics laboratory work, analyzed the data, and wrote the manuscript. RM organized sample collection and otolith ageing. ER participated in sample collection and otolith ageing. AGB and RM conceived the project. AGB, SAW, and TPO secured the funding for the epigenetic analysis. TPO provided overall supervision of the project. All authors contributed to editing the manuscript.

## Data Availability

Raw data can be found at NCBI BioProject #PRJNA1345711 and Open Science Framework at https://osf.io/jy2pb/. All data and code used to produce results, and complete analyses are available at https://github.com/emmastrand/Epigenetic_ageing. Laboratory protocols and processing can be found under the ‘protocols and lab work’ folder on the GitHub repository stated previously, in addition to Emma Strand’s GMGI Open Laboratory Notebook at https://github.com/emmastrand/GMGI_Notebook.

