## Supplemental Figures for "Building an ‘epigenetic clock’: Utilizing whole genome DNA methylation patterns to predict age in haddock, *Melanogrammus aeglefinus*"

**Additional information**

Supplemental information to include an excel spreadsheet with the following tabs:

1. Full_list: Demographics of 277 individuals sampled in Fall 2022 and Spring 2023
2. Demographic statistics: Mean, standard error, and sample size for each sex and age class determined via otolith ageing.
3. Sequence_list: 140 individuals with DNA extraction and sequence information
4. Sequencing_statistics: Run 1 and 2 sequencing coverage information
5. Bismark_statistics: DNA methylation extracted information
6. Bilsulfite_conversion: Calculations for conversion efficiency for each sample
7. Core_sites: ~3,000 CpG sites that continuously came up in generated models
8. Final_model_sites: 111 CpG loci names included in the final epigenetic ageing model
9. Age_prediction: 70/30 train-test split with otolith and epigenetic predicted ages
10. Genomic_location_summary: Number of sites in genomic regions for the clock, 100% set, and ≥90% set
11. Function_clock: Epigenetic clock CpG loci from gene regions and associated function
12. Function_90p_LargestNetwork: Functional output from STRING based on the CpG loci included in the 90% presence set.

**SUPPLEMENTAL TEXT FIGURES**

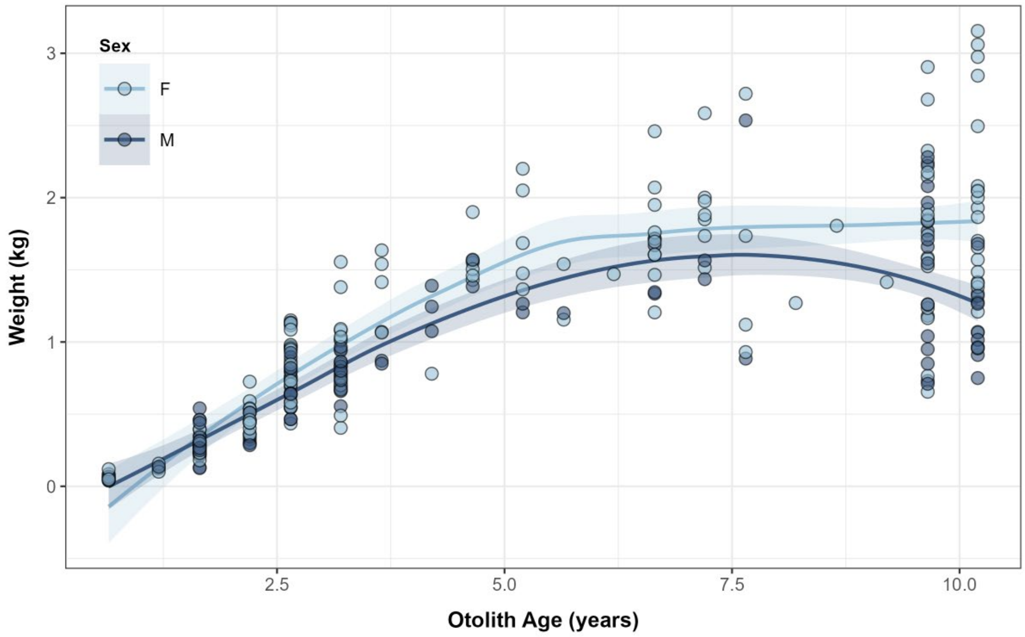


**Figure S1.** Otolith age (years) compared to fish body weight (kg) from collected individuals (n=277) during both trawl surveys. Males (dark blue) and females (light blue) are indicated by color.

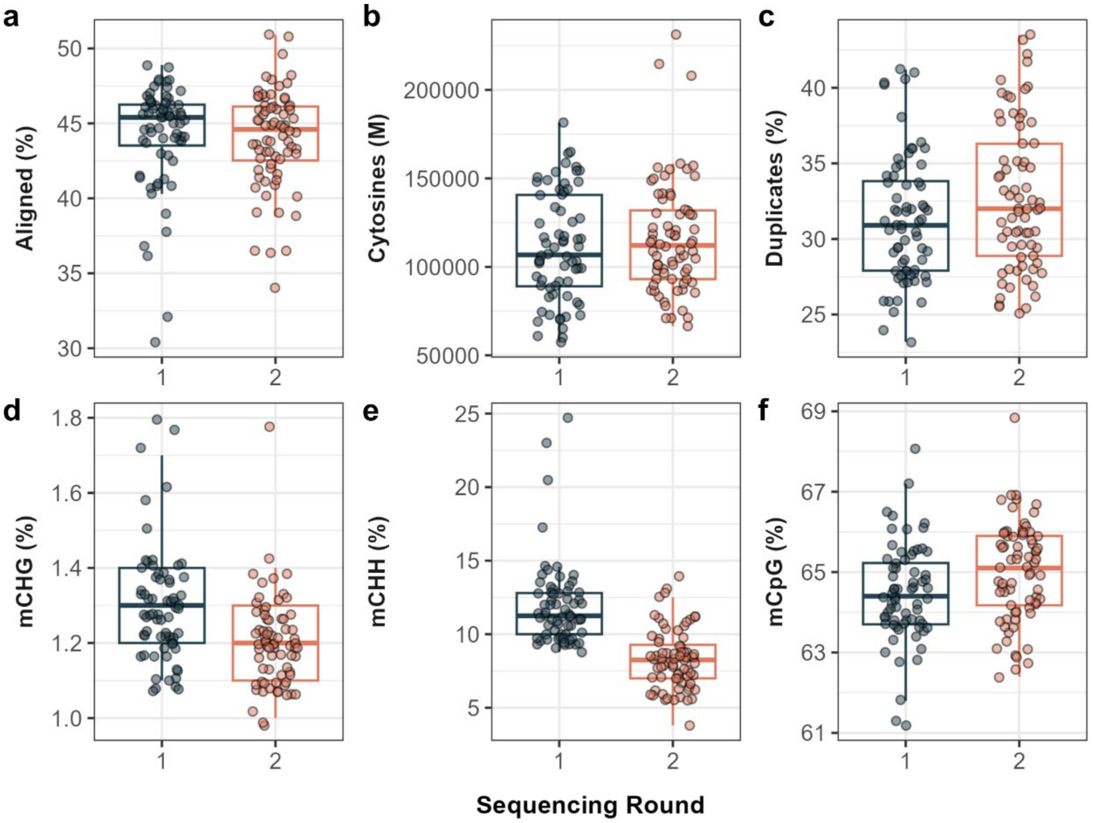


**Figure S2**. A) Percentage of reads aligned to the haddock scaffold-level genome for each of the two sequencing rounds. B) Number of cytosines (millions) with data. C) Percentage of duplicate reads. D) Percentage of methylated CHG positions across the genome. E) Percentage of methylated CHH positions across the genome. F) Percentage of methylated CpG positions across the genome. Panels D-F represent global patterns of DNA methylation.

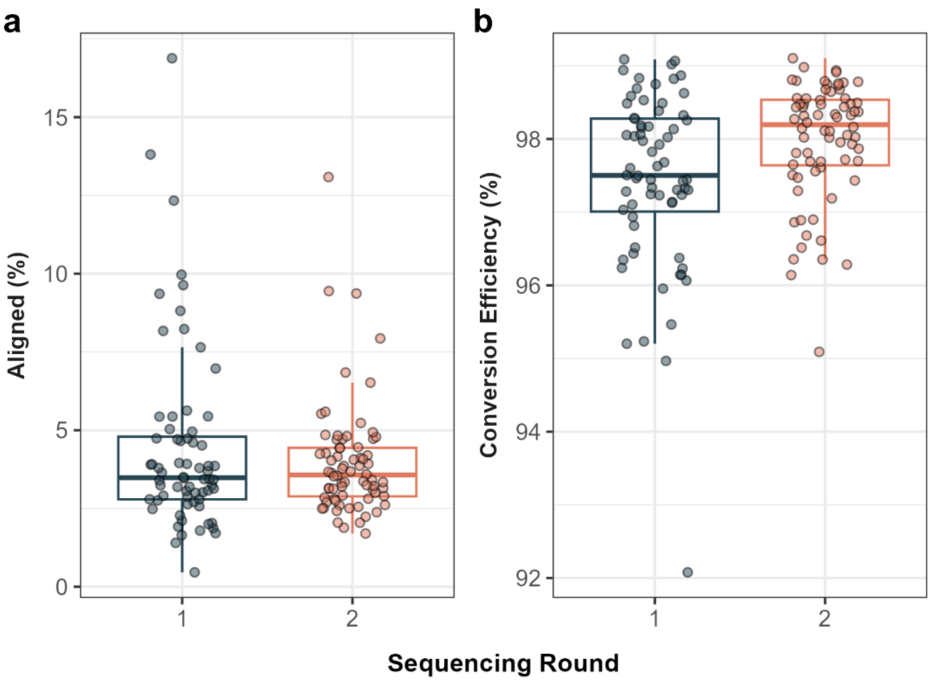


**Figure S3**. A) Percentage of reads aligned to the *E. coli* genome. B) Conversion efficiency metric (%) by run.

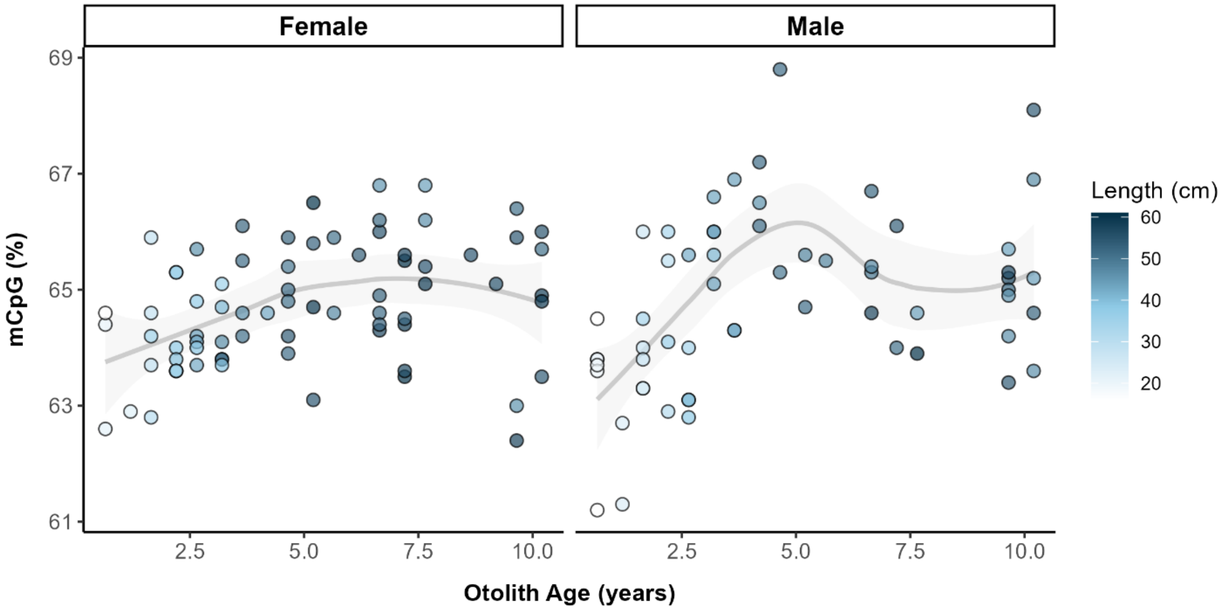

**Figure S4.** The percentage of methylated CpG positions across the genome (grey line; i.e., global DNA methylation) as a function of Otolith Age (years) split by female (left) and male (right). Each point represents an individual sample that is colored by the individual's length (cm).

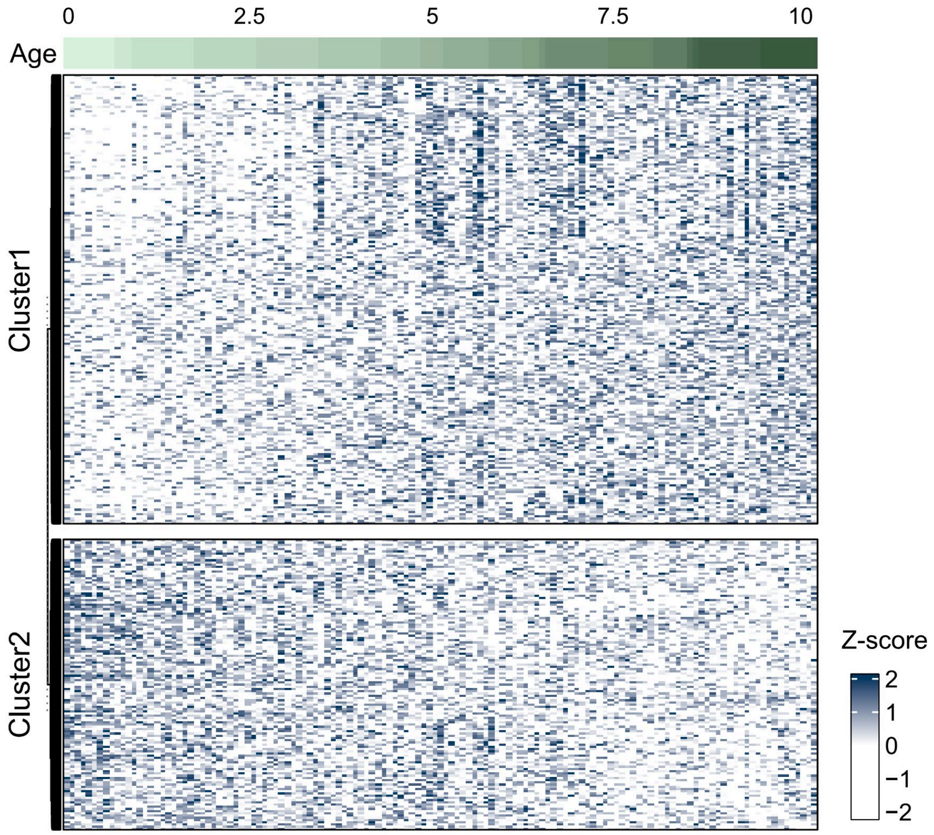


**Figure S5.** DNA methylation levels (%) were converted to a z-score and plotting by age-associated CpG loci from the ≥90% set (n=47,373; rows) across individual samples (n=140; columns). A higher z-score (darker blue) represents a higher % methylation level at that CpG loci. Older individuals are indicated by a darker green in the column color score. CpG loci cluster into 2 groups: hypermethylated with age (Cluster 1) and hypomethylated with age (Cluster 2).

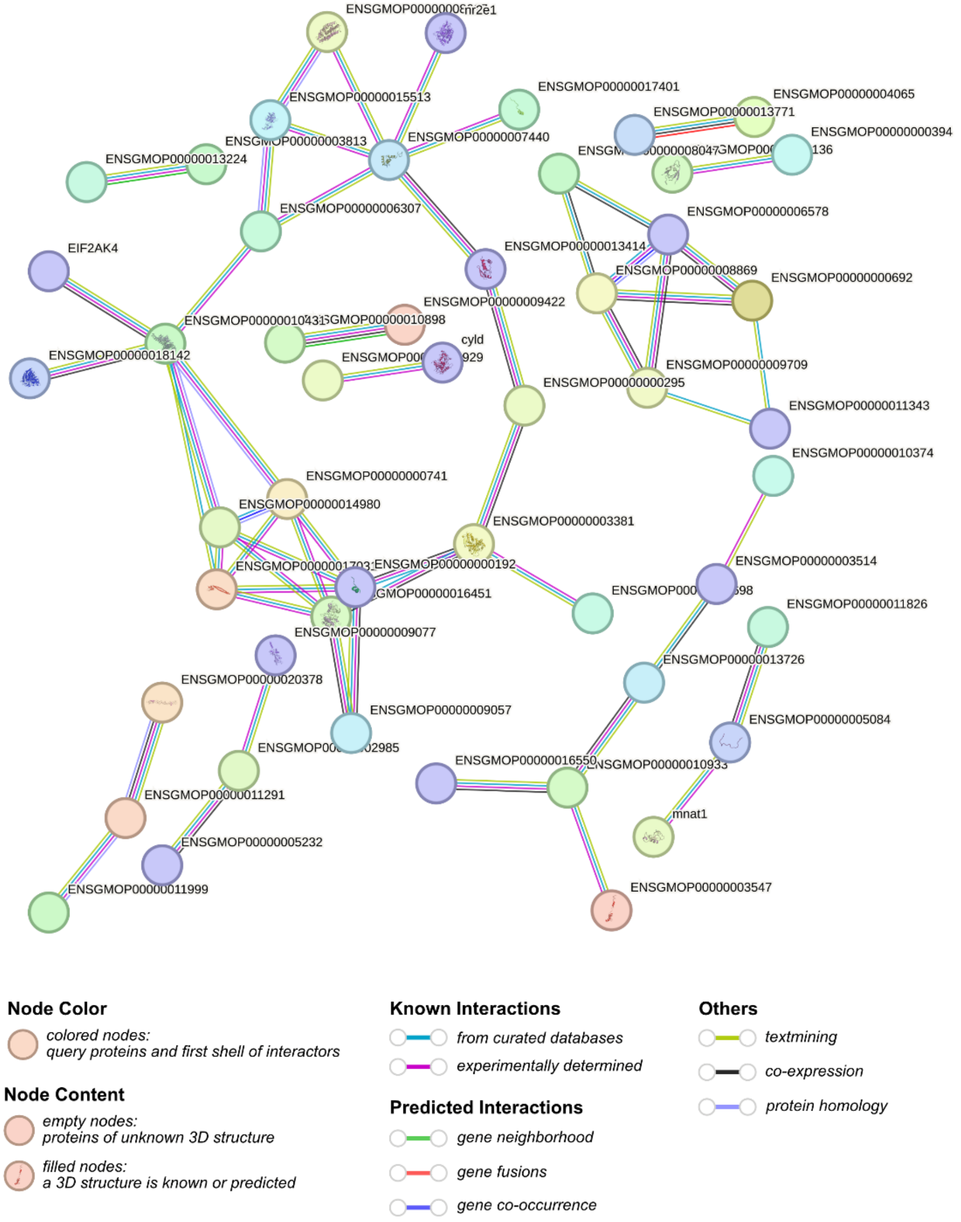


**Figure S6.** STRING network generated from the 291 protein sequences from genes with >10 age-related CpG loci found in this study. Disconnected nodes were removed from visualization and confidence level set to high confidence (0.700). Empty nodes represent nodes with unknown 3D structure and filled in nodes are those with a predicted or known 3D structure. The edge (i.e., connection) color indicates the type of interaction (e.g., co-expression data, experimentally derived data, or gene fusion data).
